# Juxta-membrane S-acylation of plant receptor-like kinases – fortuitous or functional?

**DOI:** 10.1101/693887

**Authors:** Charlotte H. Hurst, Kathryn M. Wright, Dionne Turnbull, Kerry Leslie, Susan Jones, Piers A. Hemsley

**Affiliations:** Division of Plant Science, School of Life Science, University of Dundee (at JHI), Invergowrie, Dundee DD2 5DA, UK; Cell and Molecular Science, James Hutton Institute, Invergowrie, Dundee DD2 5DA, UK; Biomedical Sciences Research Complex, University of St Andrews, St Andrews, KY16 9ST, UK

## Abstract

S-acylation is a common post-translational modification of membrane protein cysteine residues with many regulatory roles. S-acylation adjacent to transmembrane domains has been described in the literature as affecting diverse protein properties including turnover, trafficking and microdomain partitioning. However, all of these data are derived from mammalian and yeast systems. Here we examine the role of S-acylation adjacent to the transmembrane domain of the plant pathogen perceiving receptor-like kinase FLS2. Surprisingly, S-acylation of FLS2 adjacent to the transmembrane domain is not required for either FLS2 trafficking or signalling function. Expanding this analysis to the wider plant receptor-like kinase superfamily we find that S-acylation adjacent to receptor-like kinase domains is common but poorly conserved between orthologues through evolution. This suggests that S-acylation of receptor-like kinases at this site is likely the result of chance mutation leading to cysteine occurrence. As transmembrane domains followed by cysteine residues are common motifs for S-acylation to occur, and many S-acyl transferases appear to have lax substrate specificity, we propose that many receptor-like kinases are fortuitously S-acylated once chance mutation has introduced a cysteine at this site. Interestingly some receptor-like kinases show conservation of S-acylation sites between orthologues suggesting that S-acylation has come to play a role and has been positively selected for during evolution. The most notable example of this is in the ERECTA-like family where S-acylation of ERECTA adjacent to the transmembrane domain occurs in all ERECTA orthologues but not in the parental ERECTA-like clade. This suggests that ERECTA S-acylation occurred when ERECTA emerged during the evolution of angiosperms and may have contributed to the neo-functionalisation of ERECTA from ERECTA-like proteins.

## Introduction

S-acylation is a post-translational protein modification where fatty acids, such as palmitate or stearate, are covalently linked to specific cysteines through thioester linkages. No consensus sequence for S-acylation has been described and the outputs of prediction software such as CSS-Palm [1] are still some way off from being acceptably reliable. However, one structural motif has emerged; a cysteine at the cytoplasmic face of a transmembrane domain. In mammalian systems, S-acylation at cysteine residues adjacent to the transmembrane domains of single pass receptor-like portions appears to be a common feature and is important for protein functionality. Juxta-transmembrane S-acylation has been shown to affect Toll-like receptor (TLR2) trafficking to the plasma membrane and therefore regulate T-cell signalling [2]. S-acylation of the anthrax toxin receptor, TEM8, spatially segregates it in the plasma membrane away from its cognate E3 ubiquitin ligase, Cbl, to control TEM8 endocytosis and targeting to the lysosomes [3]. LRP6 is involved in the Wnt signalling pathway and requires S-acylation of the juxta-transmembrane (juxta-TM) cysteines [4]. S-acylation induces tilting of the LRP6 transmembrane span to prevent exposure of hydrophobic amino acids to the aqueous environment during trafficking through the endoplasmic reticulum. S-acylation-deficient LRP6 is recognised by the endoplasmic reticulum quality control mechanisms due to hydrophobic mismatch and targeted for ubiquitin-mediated degradation [4]. We recently identified numerous proteins from Arabidopsis as being S-acylated through a proteomics screen [5]. Included in this group of S-acylated proteins were members of the receptor-like kinase (RLK) superfamily. Receptor-like kinases are related to mammalian single pass receptor-like proteins such as TLR2 [6], and S-acylation of receptor-like kinases has been assumed to function in a similar manner to that described for single pass transmembrane proteins in mammals. However, given the evolutionary distance between plants and animals, and the differences in membrane compositions, it should not be presumed that juxta-transmembrane domain S-acylation has the same effect in both plant and animal kingdoms.

RLKs are one of the largest gene families in Arabidopsis with over 610 members [6] and their roles in a wide variety of plant processes have been well-characterised. This includes plant growth and development through the brassinosteroid hormone receptor BRI1 and its co-receptor BAK1 [7], and the control of organ formation and cell fate specification through ERECTA [8]. Other RLKs such as FERONIA and ERULUS are required for brassinosteroid-mediated cell expansion and pollen and root hair elongation [9, 10]. Additionally, wall-associated kinases (WAKs) interact with the cell wall and are required for cell expansion during plant development [11]. RLKs are also involved in abiotic stress responses through RPK1 which regulates ABA and osmotic stress signalling, and GHR1 which controls ABA- and H_2_O_2_-regulated anion channels in guard cells. Other members of the RLK superfamily are involved in plant-microbe interactions: Xa21 from *Oryza sativa* was the first RLK characterised in plants and is involved in bacterial pathogen resistance *[12]*. Other well-characterised receptors, FLS2 [13] and EFR [14] perceive the bacterial-derived peptides flagellin and elongation factor-Tu, respectively, while CERK1 and LYK4 are involved in recognition of chitinous fungal elicitors [15]. While much is known about members of the RLK superfamily, little is known about the specific role of S-acylation in RLK function.

Based on data from the original proteomics screen, we identified juxta-TM S-acylation sites in the model RLK FLS2 [5]. These data have subsequently been validated using orthogonal metabolic labelling strategies putting S-acylation of FLS2 at these sites beyond reasonable doubt [16]. Both studies demonstrated that mutation of these cysteines abolished S-acylation and we described a mild effect of their mutation on AtFLS2-mediated processes [5]. However, this work was performed with mGFP6 tagged versions of AtFLS2 expressed from the 35S promoter. We have since determined that many tags, when appended to AtFLS2, impact unpredictably on AtFLS2 function, with mGFP6 in the context of the pMDC vector series [17] used being particularly deleterious to AtFLS2 function [18]. Some of these effects can apparently be masked by dosage compensation [18] such as the 35S promoter used in our original study [5]. We therefore decided to revisit our earlier work in greater depth using untagged forms of AtFLS2 expressed close to native levels to accurately gauge the role and contribution of juxta-TM S-acylation to AtFLS2 function.

## Results

### Juxta-transmembrane domain S-acylation is not required for AtFLS2 function

We previously proposed that loss of AtFLS2 juxta-transmembrane (juxta-TM) S-acylation in AtFLS2 C^830,831^S mutants had a mild negative effect on flagellin signalling outputs when expressed as C-terminal mGFP6 fusions at 100-300x normal levels [5]. We have since determined that epitope fusions, in particular mGFP6, can have deleterious and unpredictable effects on AtFLS2 function when AtFLS2 is expressed at physiological levels, calling the biological relevance of these data into question [18]. To determine whether the minor difference in AtFLS2 function observed between AtFLS2 and AtFLS2 C^830,831^S mGFP6 tagged lines was genuine we generated *fls2/proAtFLS2:AtFLS2* [18] and *fls2/proAtFLS2:AtFLS2 C*^*830,831*^*S* lines and selected plants showing similar AtFLS2 and AtFLS2 C^830,831^S protein expression levels (Supplemental figure 1A). Following flg22 perception, AtFLS2 heterodimerises with its co-receptor BAK1 leading to the formation of an active signalling complex. This signalling complex initiates complex intracellular signalling cascades to promote plant immunity. MAPK activation is one of the earliest detectable signalling events following flg22 perception [19]. After flg22 treatment, *fls2c/AtFLS2*_*pro*_:*AtFLS2* and *fls2c/AtFLS2:AtFLS2 C*^*830,831*^*S* plant line MAPK6/3 activation was indistinguishable from Col-0, while fls2c lines showed no MAPK6/3 activation (Fig 1A). MAPK-dependent genes (NHL10, WRKY40) were similarly activated in Col-0, *fls2c/AtFLS2*_*pro*_:*AtFLS2* and *fls2c/AtFLS2:AtFLS2 C*^*830,831*^S plant lines (Fig 1B, C, Supplemental fig 1B, C). To test whether other signalling pathways were affected by loss of AtFLS2 juxta-TM S-acylation, *PHI-1*, a Ca^2+^ mediated flg22-inducible defence marker gene, was examined. Again, *PHI-1* induction was elevated in *fls2c/AtFLS2:AtFLS2 C*^*830,831*^*S* plants to a similar degree as Col-0 and *fls2c/AtFLS2*_*pro*_:*AtFLS2* control lines (Fig 1D, Supplemental fig 1D). *PR1*, a marker of salicylic acid (SA) mediated gene induction and sustained signalling, was unaffected in *fls2c/AtFLS2:AtFLS2 C*^*830,831*^*S* plant lines (Fig 1E, Supplemental figure 1E). Following longer term exposure to flg22 SA signalling is elevated and callose, a β-glucan polysaccharide, is deposited to reinforce the cell wall and prevent bacterial penetration. Callose deposition in *fls2c/AtFLS2:AtFLS2 C*^*830,831*^*S* and *fls2c/AtFLS2*_*pro*_:*AtFLS2* control lines was similar to Col-0 and in all cases much greater than in *fls2c* (Fig 1F, Supplemental fig 1F). This suggests that loss of the juxta-TM cysteines does not adversely impact any flg22-mediated AtFLS2 signalling outputs. One of the responses to prolonged flg22 exposure is the inhibition of seedling growth, and this is often used as a proxy for the overall combined signalling outputs and long-term effects of AtFLS2 activation. Growth inhibition experiments showed no deleterious effects of AtFLS2 C^830,831^S on seedlings growth inhibition responses compared to Col-0, or AtFLS2 expressing control lines (Fig 1G). This suggests that, contrary to previous conclusions from using tagged AtFLS2 C^830,831^S lines [5], juxta-transmembrane S-acylation of AtFLS2 is not required for function.

**Figure 1.**
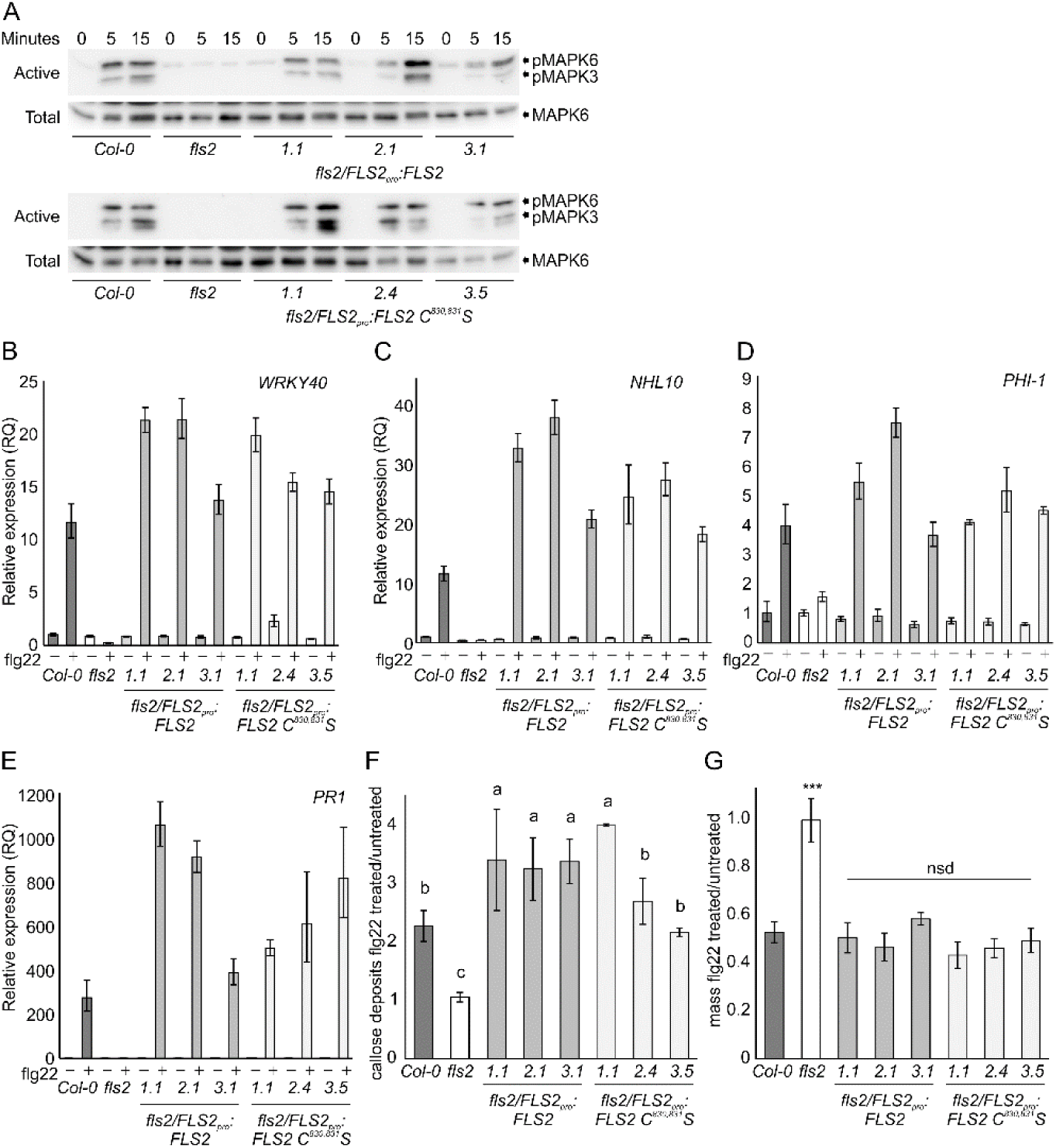
FLS2 C^830, 831^S expressing plant responses to flg22 stimulation are indistinguishable from wild type. **(A)**. MAPK activation in *fls2/FLS2pro:FLS2* and *fls2/FLS2pro:FLS2 C*^*830,831*^*S* seedlings in response to 100 nM flg22 over time as determined by immunoblot analysis. pMAPK6/pMAPK3 show levels of active form of each MAPK. MAPK6 indicates total levels of MAPK6 as a loading control. Upper shadow band in MAPK6 blot is rubisco detected non-specifically by secondary antibody. **(B)**. Induction of NHL10, **(C)** WRKY40 and **(D)** PHI-1 gene expression after 1 hour treatment with 1mM flg22 in *fls2/FLS2pro:FLS2* and *fls2/FLS2pro:FLS2 C*^*830,831*^*S* seedlings as determined by qRT-PCR. **(E)** Induction of PR1 gene expression after 24 hours treatment with 1mM flg22 in *fls2/FLS2pro:FLS2* and *fls2/FLS2pro:FLS2 C*^*830,831*^*S* seedlings as determined by qRT-PCR. Values were calculated using the DDCT method from 3 technical replicates, error bars represent RQMIN and RQMAX and constitute the acceptable error level for a 95% confidence interval according to Student’s t-test. Similar data were obtained over 2 biological repeats (additional data shown in supplemental figure 1). **(F)** Relative quantification of callose deposition following 12 hours treatment with water or 1mM flg22. Averages are shown from at least 2 biological repeats, error bars represent SEM and statistically similar groupings are denoted by a, b, c as determined by ANOVA and post-hoc Tukey-Kramer test. **(G)** Average relative seedling mass (flg22 treated/untreated) of seedlings grown in 1 µm flg22 for 10 d (14 d post germination). Data are averages of three independent biological replicates. Error bars show se. Asterisks denote statistically significant differences compared with that of the Col-0 control (***, P < 0.01) determined by one-way ANOVA and Tukey’s HSD test.

### RLK superfamily S-acylation is not confined to juxta-TM cysteines

The above findings on AtFLS2 juxta-TM S-acylation led us to question why AtFLS2 is S-acylated at these sites if, unlike reported juxta-TM S-acylation in animals [3, 4, 20], it has no discernible effect on function. In a proteomic screen for S-acylated proteins [5] we identified 16 receptor-like kinases from Arabidopsis (Supplemental Table 1) that, like AtFLS2, all contain at least one cysteine within 5 amino acids of the C-terminus of their single transmembrane domain (based on definitions from the Aramemnon database [21]). This strongly suggests that, like AtFLS2, these cysteines are likely sites of S-acylation. Analysis of the juxta-TM region of all RLK superfamily members in the Arabidopsis genome [6] (Supplemental figure S2) suggest that in total 108 contain cysteines adjacent to the TM (at or within the 5 cytoplasmic amino acids following the TM/cytoplasm interface). These cysteines represent high probability candidate sites for S-acylation [3]. The RLKs containing these cysteines are distributed widely throughout the Arabidopsis RLK superfamily (Supplemental Table 2). Two well studied receptor-like kinases that contain juxta-TM cysteines, AtLYK4 and AtERECTA, were cloned and, like AtFLS2, were found to be positive for S-acylation (Fig 2A). Using a recently devised and more sensitive S-acylation assay [22] to determine the S-acylation state of AtFLS2 expressed at native levels in Arabidopsis we observed that although S-acylation was markedly reduced in the AtFLS2 C^830,831^S mutant it was not abolished as previously described [5, 16]. This suggests that AtFLS2 contains additional or alternative S-acylation sites with low occupancy. We therefore expressed AtLYK4 and AtERECTA as EGFP fusions alongside versions containing juxta-TM cysteine mutations (AtLYK4 C^295^S, AtERECTA C^602^S) in *N. benthamiana* and assessed S-acylation state. In both cases S-acylation was still observed in the cysteine to serine mutants (Fig 2A). To verify that the juxta-TM cysteine in AtERECTA and AtLYK4 is truly a site of S-acylation, minimal constructs corresponding to AtERECTA, AtERECTA C^602^S, AtLYK4 and AtLYK4 C^295^S but lacking the kinase domain were generated. In these constructs the juxta-TM cysteines are the only cysteines remaining in the cytoplasmic portion of the protein and are therefore the only cysteines that could become S-acylated. Testing of these constructs for S-acylation confirms that the juxta-TM cysteines in AtLYK4 and AtERECTA are S-acylated (Fig 2B). All of these data combined strongly support the hypothesis that AtFLS2, AtLYK4 and AtERECTA contain S-acylation site(s) in addition to the S-acylation site adjacent to the transmembrane domain. These sites may therefore also exist within receptor-like kinases that do not contain juxta-TM cysteines. To test this we cloned FEI1, FER (Feronia), AtERL1 (AtERECTA-like 1) and BRI1 (Brassinosteroid insensitive 1) as examples of receptor-like kinases that do not contain juxta-TM cysteines and assessed them for S-acylation. Again we found that S-acylation was a feature of all RLKs tested (Fig 2C) suggesting that S-acylation of the RLK superfamily is much more common than previously proposed [5], possibly even ubiquitous, and occurs at sites other than juxta-TM cysteines.

**Figure 2.**
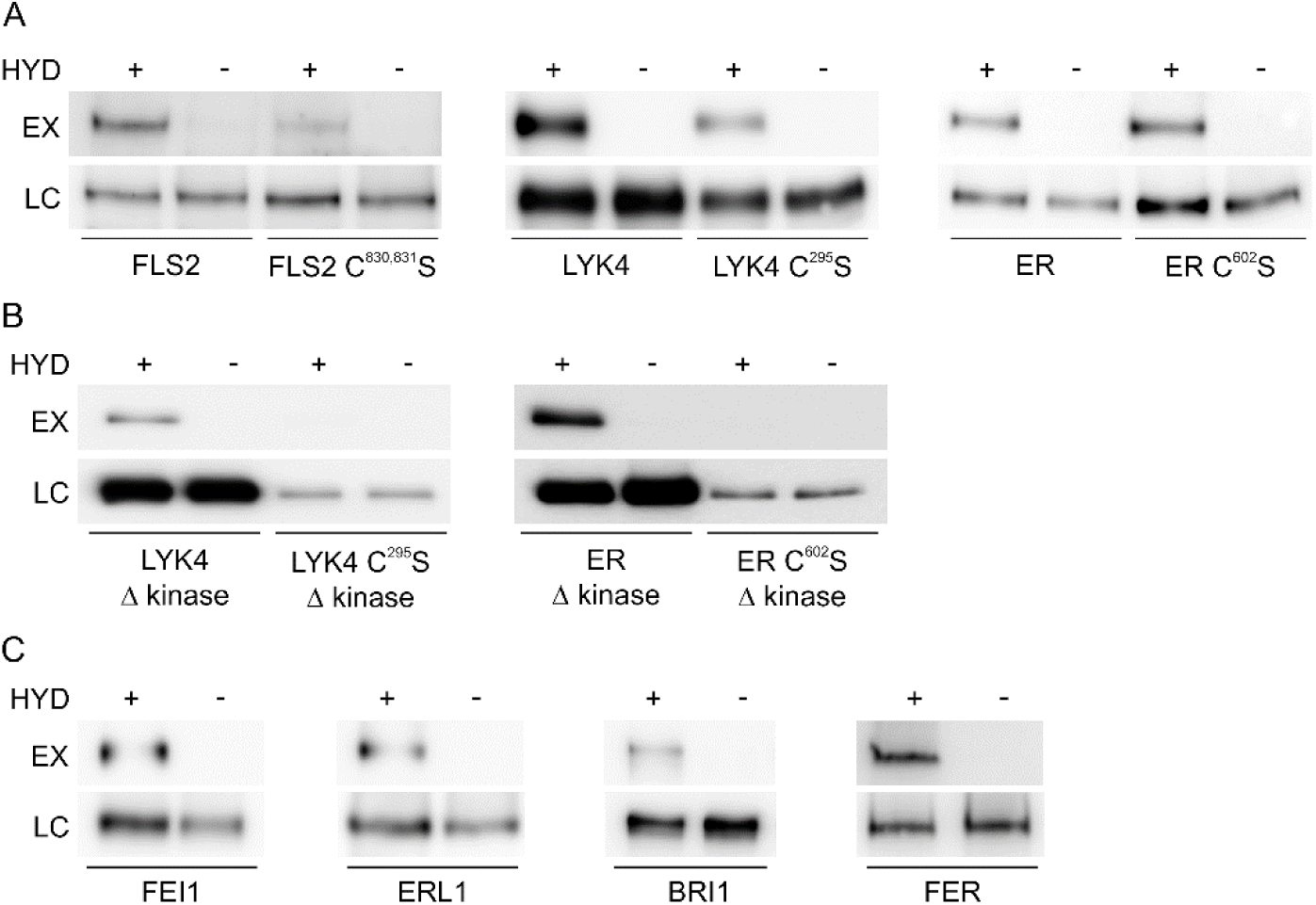
Receptor-like kinases are S-acylated at sites other than the transmembrane region. **(A)** Receptor-like kinases show reduced but readily apparent S-acylation following mutation of juxta-TM cysteine residues. S-acylation state of receptor-like kinases was determined using Acyl-biotin exchange (ABE; FLS2 variants expressed in Arabidopsis) or Acyl-RAC (LYK4 and ER variants expressed in *N. benthamiana*) methods. **(B)** Truncations of LYK4 and ER indicate that S-acylation occurs at juxta-TM cysteines similarly to FLS2. **(C)** Receptor-like inases that do not contain juxta-TM cysteines are also S-acylated. HYD indicates presence (+) or absence (-) of hydroxylamine required for S-acyl group cleavage during the thiopropyl Sepharose 6b capture (Acyl-RAC) or biotinylation step (ABE) step. Thiopropyl Sepharose 6b (Acyl-RAC) or neutravidin (ABE) enriched proteins were eluted and represent S-palmitoylated proteins (EX). Prior to thiopropyl Sepharose 6b (Acyl-RAC) or neutravidin (ABE) capture a sample was removed as an input loading control (LC).

### Juxta-transmembrane domain cysteines does not control receptor-like kinase localisation

Data from mammalian and yeast systems suggests that juxta-TM cysteine S-acylation can help direct the export of correctly folded proteins from the Golgi or ER to the plasma membrane, and prevent hydrophobic mismatch between the TM helix and the Golgi/ER membranes leading to protein turnover [4]. Our previous data suggested that AtFLS2 S-acylation was not required for PM targeting [5] and the data presented above indicate that AtFLS2 with mutated juxta-membrane S-acylation sites is fully functional and must therefore be at the plasma membrane in Arabidopsis. To determine whether the lack of effect of juxta-TM S-acylation on localisation was restricted to AtFLS2 we tested whether AtERECTA and AtLYK4 required S-acylation to traffic to the PM. Confocal imaging indicates that AtFLS2, AtLYK4, and AtER traffic to the PM in the absence of juxta-TM cysteines, with the intracellular distribution of the juxta-TM mutant forms being indistinguishable from wild type (figure 3).

**Figure 3.**
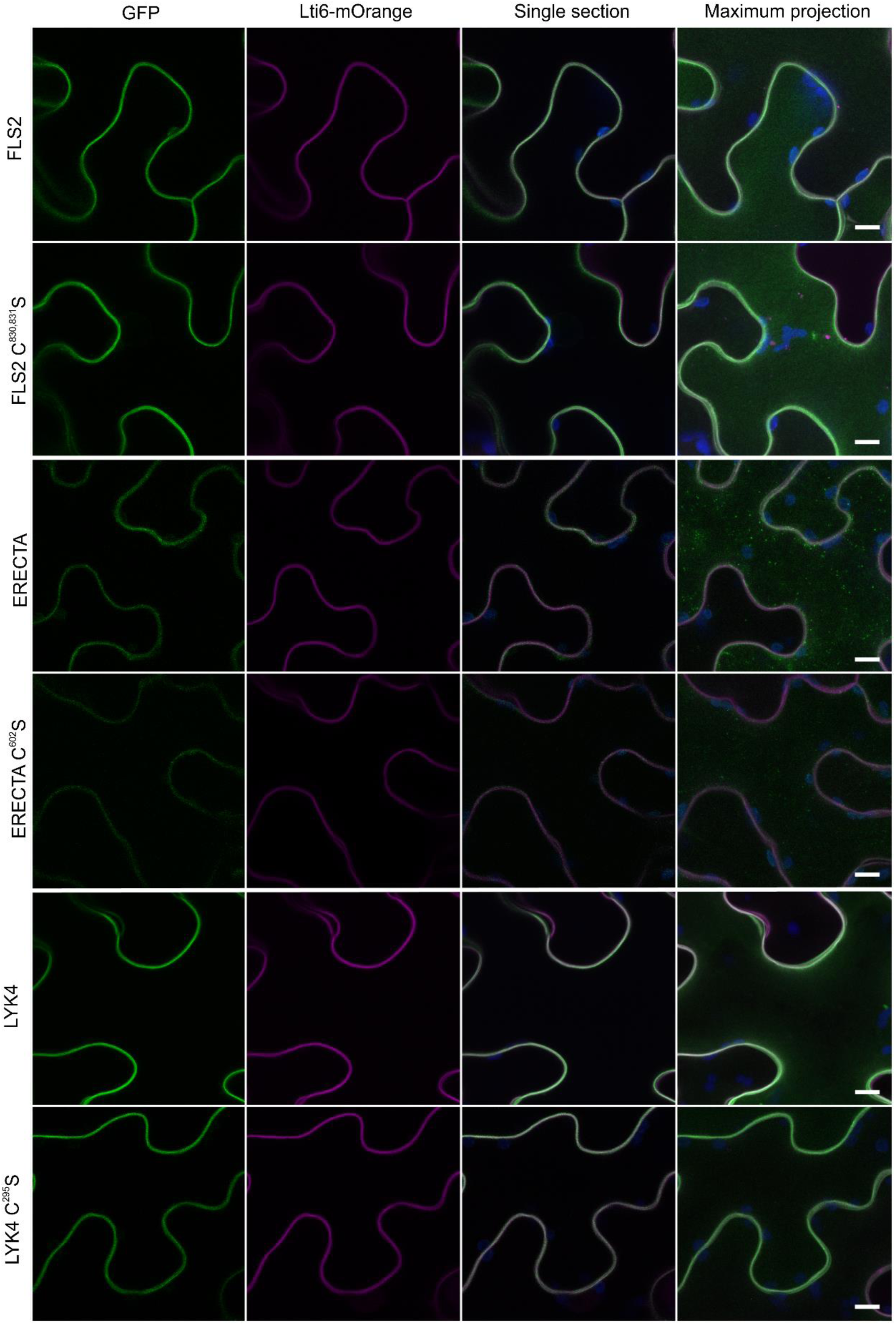
Receptor-like kinase localisation to the plasma membrane does not require S-acylation. Receptor-like kinase variants fused to EGFP (green) were imaged in mOrange-Lti6b (magenta) expressing plants. Chlorophyll auto-fluorescence is shown in blue. Scale bar represents 10 μm.

### Juxta-membrane S-acylation in not necessarily conserved between orthologues

If juxta-TM S-acylation were major or essential determinants of a receptor-like kinases’ function it would be expected that juxta-TM sites of S-acylation would be under strong selective pressure over evolutionary time. As a result, clear patterns of gain or loss of S-acylation would be expected with the presence/absence of juxta-TM S-acylation being highly conserved between RLK orthologues. If juxta-TM cysteines occur through random mutation and confer no particular benefit, then they will be under neutral selection and their presence would be more or less random with no clear evolutionary pattern. Using the Aramemnon database of angiosperm membrane proteins we examined orthologues of Arabidopsis RLKs containing juxta-TM cysteines found both through proteomics or tested here for S-acylation, and examined them for juxta-TM cysteines. As can be seen in supplemental figure 4, using the previously defined evolutionary receptor-like kinase sub-family groupings [6], there is no particular overall pattern of juxta-TM cysteine occurrence either between or within receptor-like kinase sub-families across different species. However, some orthologues, such as those for ERECTA, TMK3 or THESEUS, do appear to show conservation of juxta-TM cysteines across species. Interestingly the closest relatives of ERECTA, ERECTA-like1 and 2 (ERL1/2), do not have a juxta-TM cysteine in any species examined. This suggests that the juxta-TM cysteine of ERECTA appeared at the point when ERECTA diverged from the ERL1/2 clade, and has since been maintained in all extant angiosperm lineages. Given these data, and how S-acylation occurs sporadically throughout the Arabidopsis receptor-like kinase family tree (Supplemental figure 3), we propose that receptor-like kinase juxta-TM S-acylation has arisen and been lost multiple times over plant evolution and receptor-like kinase family diversification. Based on our data we propose that many juxta-TM cysteine-containing receptor-like kinases, such as AtFLS2, where orthologue juxta-TM cysteines are not conserved, may be fortuitously S-acylated but that S-acylation may become a fixed feature if it provides a novel beneficial function.

## Discussion

Our previous work on S-acylation of AtFLS2 suggested that loss of S-acylation had a very mild effect on AtFLS2 function [5]. Since performing this work we have discovered that different epitope-tagged forms of AtFLS2 behave in an unpredictable manner, ranging from essentially fully functional to almost inactive [18]. The tag used in our original study [5], mGFP6, has a marked effect on AtFLS2 function when expressed at levels similar to endogenous AtFLS2 [18] but these effects can be masked by overexpression [5]. In light of this work, we reassessed our previous data using untagged forms of AtFLS2 lacking juxta-TM S-acylation, expressed at levels similar to endogenous AtFLS2. Using these lines we found that loss of AtFLS2 juxta-TM S-acylation had no discernible effect on any FLS2 mediated function or output tested. This surprising result led us to conclude that AtFLS2 juxta-TM S-acylation is likely fortuitous. This stands in stark contrast to the many documented examples of juxta-TM S-acylation being required for the trafficking, function or regulation of diverse single-pass transmembrane receptors in animals [2–4, 20].

S-acylation does not appear to be specified only by primary sequence; rather secondary and tertiary structure also likely govern whether S-acylation occurs. Common structural motifs for S-acylation include transmembrane domains followed by one or more cytosolic cysteine residues. Cysteines adjacent to a transmembrane domain likely evolved as an S-acylation motif as a result of the active site of S-acyl transferases being located at the membrane-cytosol interface, placing active site and S-acyl accepting cysteine in the same plane [23]. Juxta-TM cysteines are likely to arise comparatively often by chance as 13/61 amino acid encoding codons (~21%) can alter to code for cysteine with only one base change. Furthermore, the amino acids encoded by these 13 codons (Phe, Tyr, Gly, Ser, Arg, Trp) are over-represented in α-helical transmembrane domains and junctions between transmembrane domains and the aqueous environment [24]; the very region where S-acylated cysteines are found in receptor-like kinases. We propose that chance mutation may frequently result in cysteines occurring adjacent to transmembrane domains where they can be readily S-acylated by S-acyl transferases that recognise the “TM and juxta-TM cysteine” motif. Recognition of these new substrates by S-acyl transferases is eminently possible due to the common occurrence of this motif in pre-existing S-acylated proteins [20] and the rather lax specificity of many S-acyl transferases [25, 26]. Due to their reactivity, cysteine residues are not commonly found as surface exposed residues and tend to be under negative selection pressure unless they are required for a specific biological function [27]. However, S-acylation would essentially cap the cysteine residue and prevent reaction with other cellular constituents. As long as S-acylation does not negatively impact upon receptor-like kinase function the cysteine will therefore persist under neutral selection. We propose that S-acylation of AtFLS2 at Cys830 and 831 falls into this category as they have no apparent effect on function and no AtFLS2 orthologue examined outside of the *Brassicaceae* possesses these sites, and absolute sequence and positional conservation within the *Brassicacea* is poor. This also seems to be the case for LYK4 as S-acylation of LYK4 orthologues is not conserved. For instance, examination of the four dicot and three monocot species presented in supplemental figure 4 indicates that LYK4 orthologue S-acylation either arose in the last common ancestor of eudicots and monocots and was subsequently lost in *P. trichocarpa, S. lycopersicon* and *Z. mays* lineages or has arisen multiple times independently (Arabidopsis, *C. melo, O. sativa* and *B. distachyon*). If, however, S-acylation imparts a beneficial function or character to the receptor-like kinase, then it will be positively selected for and maintained over evolutionary time and distance. We hypothesise that this is the case for ERECTA. The ERECTA-like family of receptor-like kinases is present in all embryophytes, and gave rise to the ERECTA sub-clade [28] following the epsilon whole genome duplication event in the last common ancestor of all extant Angiosperms [29]. Interestingly, all ERECTA orthologues that we have examined (supplemental figure 3) contain positionally conserved juxta-TM cysteines suggesting that, from its point of origination, ERECTA was S-acylated. This is further supported by examination of ER and ERL orthologues from *Amborella trichocarpa*, considered to be the sister group to all other extant angiosperms [30], that follow the same pattern. We were unable to find evidence of ERECTA-like clade juxta-TM cysteines encoded in any gymnosperm genome. This would suggest that ERECTA S-acylation, unlike in AtFLS2, serves a purpose and we hypothesise that S-acylation enabled and characterised the emergence and neo-functionalisation of ERECTA from the ERECTA-like clade in a common ancestor of all extant sequenced angiosperms. We are currently investigating this intriguing role of S-acylation as a potential evolutionary driver in ERECTA function. This work has also revealed that all receptor-like kinases appear to be S-acylated at sites independent of any juxta-TM cysteines. This exciting finding indicates that S-acylation of receptor-like kinases is a general feature of the superfamily rather than a curious feature of a minority although the identity and role of these additional sites has yet to be determined.

## Material and methods

### Plant lines and growth conditions

All Arabidopsis thaliana lines used in the work are Col-0 ecotype. *fls2* (SAIL_691C4, [31]) and *fls2c*/AtFLS2_pro_:AtFLS2 [18] plant lines have been previously described. Plant material for Arabidopsis experiments was grown on 0.5x Murashige & Skoog (MS) medium containing 0.8% (w/v) agar. Seeds were stratified for 48h at 4°C and germinated under a 16/8 h light/dark cycle at 20°C in MLR-350 growth chamber (Panasonic). *Nicotiana benthamiana* plants expressing mOrange-LTI6b fluorescent protein as a plasma membrane marker (mOrg-LTI; [32]) were used for confocal imaging. Transgenic and wild-type *Nicotiana benthamiana* plants were grown in commercial compost and maintained under glasshouse conditions of 16/8 h light/ dark cycle, daytime temperature 26°C maximum, night-time 22°C for 4-5 weeks before use.

### Constructs

AtFLS2_pro_:AtFLS2 C^830,831^S (pK7WG0, [33]) constructs were created using the same promoter and open reading frame as found in *fls2/AtFLS2*_*pro*_:*AtFLS2* described previously [18]. Cysteine codon mutants and reading frame truncations were generated using Q5 site-directed mutagenesis (NEB). Transgenic Arabidopsis plant lines were generated by *Agrobacterium tumefaciens*-mediated floral dip transformation [34] and selected for homozygosity at T3. Additional receptor-like kinases were cloned from genomic Col-0 DNA without a stop codon into pENTR D-TOPO before sequencing and recombination into pK7FWG2 [33] or pGWB17 [35] for expression from the 35S promoter as C-terminal EGFP or 4xMYC fusions respectively.

### Transient expression in N. benthamiana

Leaves of 4-5 week old wild type and mOrg-LTI [32] expressing *Nicotiana benthamiana* plants were syringe infiltrated with *Agrobacterium tumefaciens* GV3101 pMP90 containing expression plasmids for each receptor-like kinase alongside p19, each at an OD_600_ of 0.1. Plants were maintained under standard growth conditions for 48-60h before use in experiments.

### Gene expression analysis

Gene expression level analysis was carried out as previously described [18, 36]. Gene expression was analysed by quantitative reverse transcription PCR. 10 seedlings of each genotype 10 days post-germination were treated with 1μM flg22 for the indicated times in 2 mL 0.5x MS liquid medium. RNA was extracted using an RNeasy mini kit with on-column DNAse digestion according to manufacturer’s instructions (QIAgen). 2 μg RNA was reverse transcribed using a high capacity cDNA reverse transcription kit (Applied Biosystems). Expression levels were normalised against PEX4 (At5G25760, [37]). Relative quantification was calculated using the ΔΔC_T_ method [38]. Error bars represent RQ_MAX_ and RQ_MIN_ from three technical replicates and constitute a 95% confidence interval, according to a Student’s t-test.

### MAPK activation

MAPK activation assays were performed as described previously [39]. Six seedlings of each genotype 10 days post-germination were transferred to 12-well plates containing 2 mL 0.5× MS liquid media for 24 h and then treated with 100 nM flg22 for the indicated times in 2 mL 0.5× MS medium. 30μg total protein was loaded per sample.

### Immunoblot analysis of protein levels

Proteins were extracted from pools of seedlings as previously described [22] separated by SDS-PAGE and transferred to PVDF membrane using wet transfer methods and blotted for active and total MAPK [39] or AtFLS2 [22]. Antibodies used were as follows: α-p42/44 MAPK: CST (#9102); α-MPK6: Sigma-Aldrich (A7104); α-EGFP: Roche (11867423001); α-MYC: Thermo (MA1-21316).

### Callose staining

Callose staining was carried out as previously described [40]. Seedlings were destained overnight before aniline blue stain was applied for 3h in the dark. Callose images were obtained from cotyledons 10 days post-germination treated for 12h with 1 μm flg22 or water. Cotyledons were mounted in 50% glycerol and imaged using a LEICA DMLFS microscope with a 5x objective lens and UV D filter. Callose was quantified using an image processing macro in FIJI [41] adapted from [42]. Colour images were converted to 8-bit greyscale before analysis.

### Seedling growth inhibition

Seedlings growth inhibitions were carried out as previously described [18, 43]. Four days post-germination, 10 seedlings of each genotype were transferred to 12-well plates containing 2 mL 0.5× MS liquid media with or without 1 μM flg22, ensuring cotyledons were not submerged. Seedlings were grown for a further 10 days, blotted dry and the fresh weight of pooled seedlings in each genotype for each treatment weighed and an average calculated.

### S-acylation assay

S-acylation assays were carried out as previously described [22]. Briefly samples were ground to a fine powder in liquid nitrogen and resuspended in 2 mL lysis buffer (50mM Tris HCl pH 7.2, 5 mM EDTA, 150 mM NaCl, 2.5% SDS, 10 mM N-ethylmaleimide, 5 μl.ml^−1^ protease inhibitor [P9599, Sigma-Aldrich], 100mg PVPP). Samples were mixed gently for 10 minutes at room temperature and centrifuged at 16,000 xg for 1 minute. The supernatant was retained and total protein concentration was determined by BCA assay (ThermoFisher). 1 mg of protein was diluted to a final volume of 1 mL with lysis buffer (no PVPP) and incubated for 1h at room temperature with gentle mixing. 12μl 2,3-dimethyl 1,3-butadiene was added to each sample and incubated at 25°C in a shaking incubator for 1h to remove unreacted NEM. 1/10^th^ volume chloroform was added to each sample, briefly vortexed and centrifuged to achieve phase partitioning. The upper phase was retained split in two 0.5mL samples; one treated with 1 M hydroxylamine pH 7.2 and the other with an equal volume of water. 10 μl loading control was then taken from each sample and incubated at RT for 1 hour before being combined with 10 μl 2x reducing SDS-PAGE loading buffer. 20 μl equivalent of a 50% slurry of thiopropyl Sepharose 6b beads pre-swollen in lysis buffer (no NEM) was added to remaining samples and samples incubated for 1h with gentle mixing for 1 h at room temperature. Beads were washed three times in lysis buffer (no NEM) for five minutes each and the supernatant aspirated after the final wash. Proteins were eluted in 25 μl 2x reducing SDS-PAGE loading buffer at 37°C for 30 minutes with agitation.

### Confocal imaging

Cells were imaged using a Nikon A1R confocal laser scanning microscope mounted on an NiE upright microscope fitted with an NIR Apo 40x 0.8W water dipping lens and GaASP detectors. Images are false colour single confocal sections or maximum projections of Z-stacks produced using NIS-elements AR software. GFP was excited at 488 nm with emissions gathered at 500-530 nm with simultaneous collection of chlorophyll fluorescence at 663-738nm. mOrange was excited sequentially at 561 nm with emissions gathered at 570-620 nm.

### Bioinfomatic analyses

Biomart [44] was used to query EnsemblPlants [45] and extract the sequences of the 609 Arabidopsis RLK superfamily proteins defined in the work of Shiu and Bleecker [6]. Custom Perl scripts were used to extract sequences of the TM domain as defined in the ARAMEMNON database [21] plus 10 residues after the end of the TM domain. Custom scripts were then used to identify those proteins with cysteines within the region −1,+5 of the end of the TM domain. Figures were generated using custom scripts and ESPript3 [46].

### Accession numbers

Accession numbers are as follows: FLS2, At5g46330; ER, At2g26330; LYK4, At2g23770; FER, At3g51550; BRI1, At4g39400; FEI1, At1g31420; ERL1, At5g62230; NHL10, At2g35980; WRKY40,; PHI-1, At1g35140; PR1, At2g14610; PEX4, At5g25760; MPK6, At2g43790; and MPK3, At3g45640.

## Acknowledgements

We would like to thank Dr. Nana Keinath and Profs. Antje Heese, Silke Robatzek and Cyril Zipfel for helpful discussions during the preparation of this manuscript. This work was funded by UK Biotechnology and Biological Sciences Research Council grants BB/M024911/1 and BB/M010996/1 to P.A.H.

## Author contributions

PAH conceived and developed the work. CHH, DT, KMW, SJ, KL and PAH performed the experiments and analysed the results. PAH and CHH wrote the manuscript with input from DT, KMW, SJ and KL.

