## Supplemental figure 1 for "Juxta-membrane S-acylation of plant receptor-like kinases – fortuitous or functional?"

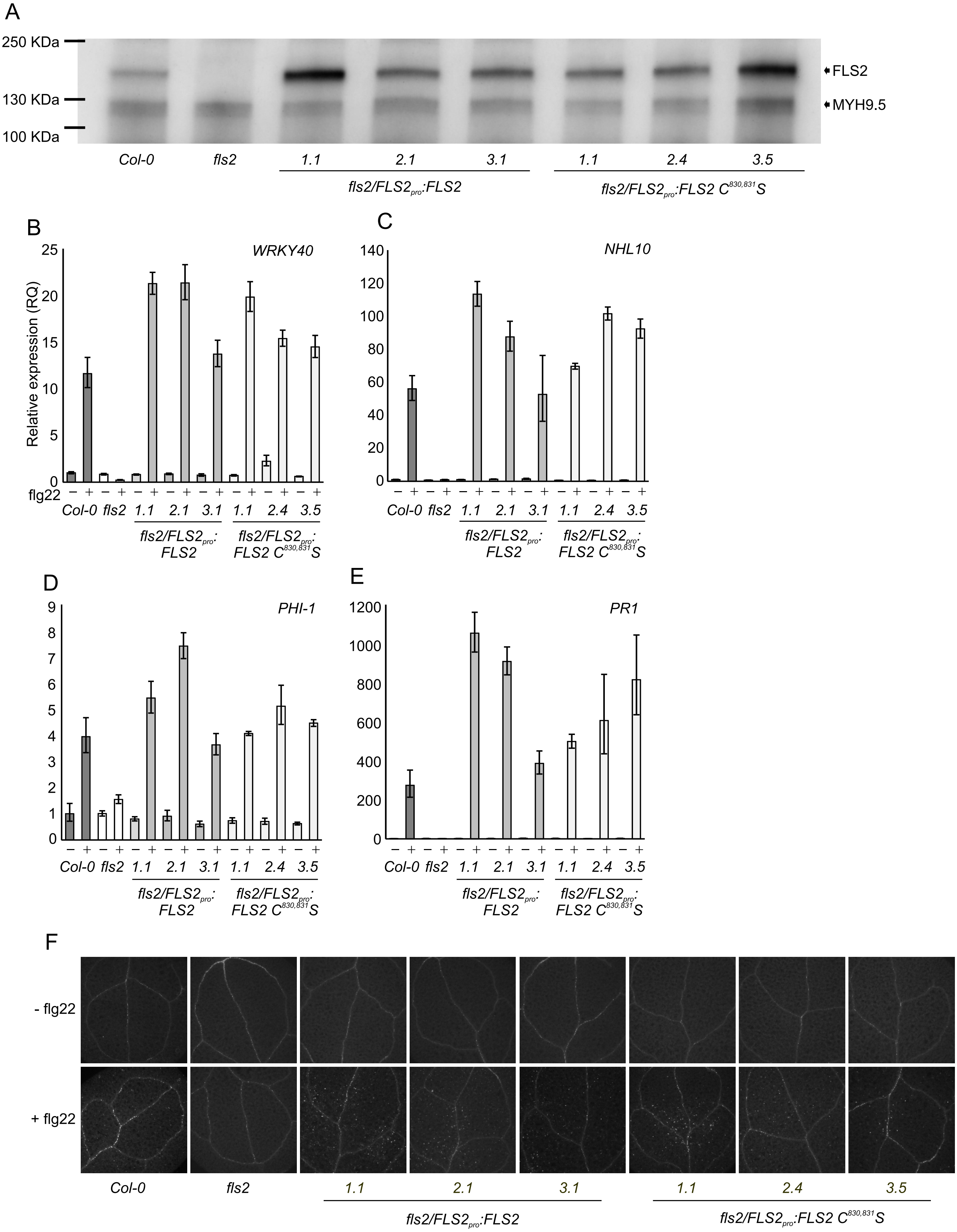


**Supplemental figure 1. FLS2 $C^{830,\ 831}$S expressing plant responses to flg22 stimulation are indistinguishable from wild type (A).** FLS2 protein expression in *fls2/FLS2pro:FLS2 and fls2/FLS2pro:FLS2 $C^{830,831}$S* seedlings in transgenic lines used in this study. Lower band labelled MYH9.5 is also recognised by the antibody used and serves as a loading control. **(B).** Induction of *WRKY40*, **(C)** *NHL10* and **(D)** *PHI-1* gene expression after 1 hour treatment with 1μM flg22 in *fls2/FLS2pro:FLS2 and fls2/FLS2pro:FLS2 $C^{830,831}$S* seedlings as determined by qRT-PCR. **(E)** Induction of *PR1* gene expression after 24 hours treatment with 1μM flg22 in *fls2/FLS2pro:FLS2 and fls2/FLS2pro:FLS2 $C^{830,831}$S* seedlings as determined by qRT-PCR. Values were calculated using the $\Delta\Delta C_T$ method from 3 technical replicates, error bars represent RQMIN and RQMAX and constitute the acceptable error level for a 95% confidence interval according to Student's t-test. Similar data were obtained over 2 biological repeats (additional data shown in figure 1). **(F)** Representative images of callose deposition following 12 hours treatment with water or 1μM flg22 used to generate graph in figure 1F.
