## Supplemental figure 2 for "Juxta-membrane S-acylation of plant receptor-like kinases – fortuitous or functional?"

```
                     1        10        20        30
AT1G01540            LWVVIGILLGSLIVIALFLLSLCLTSRRKNR
AT1G07560            PVVASLASLAAIIAMIALLFVCIKRRSSSRK
AT1G11050            ALALGITGAIFGALVIAGLICLYFRFGKAVK
AT1G11130_SUB        QRIILVVSSVAIIVLVSGLCVTLWRCCRSKI
AT1G11330_RDA2[P]    LAVMIAAPVIGVMLIAAVCVLLACRKYKKRP
AT1G11410            RLVLILISLIAVVMLLLISFHCYLRKRRQRT
AT1G16260            AILSGTLSSGLLLLIFGMWLLCKANRKRKVA
AT1G17230_EXL5       KILTITCIVIGSVFLITFLGLCWTIKRREPA
AT1G17750_PEPR2      ALIAAGSSLSVLALLFALFLVLCRCKRGTKT
AT1G21250_WAK1       WTTILLVTTIGFLVILLGVACIQQRMKHLKD
AT1G25320[P]         AIVAIVVCDFIGICIVGFLFSCCYLKICARR
AT1G26150_PERK10     AAVVGVSIGVALVLLTLIGVVVCCLKKRKKR
AT1G29740            LILGIAALIVSLSFLILGALYWRICVSNADG
AT1G29750_RKF1       AYIAIGIGAPCLIIFILGFLWICGCLPRCGR
AT1G30570            WISVGAGIAIIIFFVFLGILVVCLCKKRRSK
AT1G34300            HLWIVAVAVIAGLLGLVAVEIGLWWCCCRKN
AT1G35710            LVVWILVPILGVLVILSICANTFTYCIRKRK
AT1G48480_RKL1[P]    IAGIVIGCVVGFALIVLILMVLCRKKSNKRS
AT1G49270_PERK7      GAVIGLVAAAGILFIVMILLCVCCFRKKKKK
AT1G50610_PRK5       YTLAIILIVIGIILVIIALVFCFVQSRRRNF
AT1G51810            TIPIVASIGSVVAFTVALMIFCVVRKNNPSN
AT1G51910            LVAILASVAGVIALLAIFTICVIFKREKQGS
AT1G51940_LYK3       IWIVGGLGVVLALLVLCILVCICLRSSSCSS
AT1G56720            VWEVIGIAVALLIIAILSVLSCCLTSKKKSR
AT1G60630            IISGSICGGILILLLTFLLICLLWRRKRSKS
AT1G61360            IITVATLSLSVCLILVLVACGCWRYRVKQNG
AT1G61430            TIVASTVSLTLFVIFGFAAFGFWRCRVEHNA
AT1G61480            IIVASIVSLSLFVILAFAAFCFLRYKVKHTV
AT1G61500            IVASIVSLTLFMILGFTAFGVWRCRVEHIAH
AT1G64210            LAFLLILSAACVLCVSGLSFIMITCFGKTRI
AT1G66150_TMK1       IGIIVGSVLGGLLSIFLIGLLVFCWYKKRQK
AT1G66880            AGIAVASVSGLAILLLAGLFLCIRRRRKTQD
AT1G66910            IVIYVLGIGAASFAMMGVILVVTCLNCLIRR
AT1G66930            AIIIGIFVALCTIGGFIAFLVLLCPCCKVRI
AT1G66980            IRIVSWSVAGVVLFLVLLTLVFCFHRKRETR
AT1G67000            YTFIALGALTGVVIVFLVLLCPCFRVQIFRK
AT1G67720            GISIAAVAILLLLVGGSLVLLCALRKTKRAD
AT1G68400            SLIAIILGDFIILSFVSLLLYYCFWRQYAVN
AT1G70520_CRK2       VVIVVSVLSSVVVFMIGVAVSVYICKRRTIK
AT1G73080_PEPR1      IVLIAVLSSLLVLVVVLALVFICLRRRKGRP
AT2G01820[P]         KIIVPVVGGVVGALCLVGLGVCLYAKKRKRP
AT2G16250            LAAVGGGVAFILLFVILPIILVLCMRHRRRA
AT2G18470_PERK4      NTAIIVGVLVGAGLLMIVLIIVCLRRKKKRK
AT2G20850_LRR-RLK    IIWISILGAFSFVVLALVCLLCGRKCLRKRE
AT2G23450_WAKL14     TIVGGTVGGAFLLAALAFFFFCKRRRSTPLR
AT2G23770_LYK4[*]    YALAGVLGGALVLSVIGAAIFCLSKKKTKTQ
AT2G24230            LALAVTLSTMCLLIGALIFVAFGCRRKTKSG
AT2G26330_ER[*]      RAAILGIAIGGLVILLMVLIAACRPHNPPPF
AT2G26730[P]         IVAIIVASALVALLLLALLLFLCLRKRRGSN
AT2G28250_NCRK       KLVIVILLFCGVLISLAFLASMICYICRKDK
AT2G29250_LECRK32    TMLIIIIAASAIFGILILSFLAVCFFRRTEN
AT2G30940            LITASASIAFLLVLIISVLLCFIFHRRRCSQ
AT2G33580LYK5[P]     WIYIGIGIGAGLLLLLSILALCFYKRRSKKK
AT2G36570_PXC1       GIIAAVIGGCVAVIVLVSFGFAFCCGRLDRN
AT2G37050            GVIIGASVGAFVLLIATIISCIVMCKSKKNN
AT2G39360            GLIAGLSAALCVALVFGVVVSWWCIRKRRRR
AT2G48010_RKF3[P]    VLVSSFSVLLVASVLVITAWFWYCRRKKSKL
AT3G02880_KIN7[P]    AIVGIVIGCVVGLLLLLLILFCLCRKRKKEE
AT3G08680[P]         GAIVGIAVGGSVLLFIILAIITLCCAKKRDG
AT3G14840_LIK1[P]    TVVGSVIASTVFLVLLIGGILWWRGCLRPKS
AT3G18810_PERK6      AIVGIVAGAGLLFLVMILFCVCCCRKKKKKH
AT3G20190_PRK4       FFIIAIVLIVIGIILMIISLVVCILHTRRRK
AT3G24540            TGAVVGISIGGGVFVLTLIFFLCKKKRPRDD
AT3G24550_PERK1[P]   GVVVGIAIGGVAILVILTLICLLCKKKRRRR
AT3G46290_HERK1      LGLIVGSAIGSLLAVVFLGSCFVLYKKRKRG
AT3G47110            ITICVSAVMAALLLLCLCVVYLCWYKLRVKS
AT3G47580            AILVSIGIALLLLLVIASMVLCWFRKRRKNQ
AT3G51740_IMK2[P]    ILIAIGALLAILLLLCCILLCCLIKKRAALK
AT3G55950_CCR3       LLAFAIVGSVGAFAGICSVVYCLWTGVCLGK
AT3G58690            ALVAIVVLACLALSSLFVAFSYYCYIRNKVS
AT4G01330            LWVVIGILLGSLIVIALFFLSLCLTSRRRNR
AT4G02010            NLILIFSIAAGVLILAIITVLVICSRALREE
AT4G08850_MIK2       IYILVPIIGAIIILSVCAGIFICFRKRTKQI
AT4G11460_CRK30      TIIGIVIVVAMVIIMALLALGVSVCRSRKKY
AT4G18640_MDIS2      WLYVVIAVASFVGLLIIVAVIFFCRKRAVKS
AT4G22130_LRR-RLK    VVTGIVFGSLFVAGIIALVLYLCLHKKKRKV
AT4G23150_CRK7       NVVVVAVVVPIIVAVLIFIAGYCFFAKRAKK
AT4G23180_CRK10      VLVIAIVVPIIVAVLLFIAGYCFLTRRARKS
AT4G23250_CRK17      GTIAAIVVVVVVTIILIVVGLVICKRRKQKQ
AT4G23300_CRK22      VVVITVPAVVIVLILVVLGFFICWRRKSLQR
AT4G23740            VFLLIVIAVSIVVITALAFVLTVCYVRRKLR
AT4G25390            FPPFVVAGAGAGFSLFITLSVCFCKFSRKRS
AT4G27300_SD11       VVGMVVGSVVAIAVVLVVVFACFRKKIMKRY
AT4G28490_RLK5       VWILLTIFLLAGLVFVVGIVMFIAKCRKLRA
AT4G28670_CRK43      LTYIFVISMVGVLAIAAGFWCGKCFYMRTSP
AT4G29180            MVPIVVSTLVIILIAALAIICIMRRESKIMY
AT4G30520            LAIALSVSLGSVVILVLALGSFCWYRKKQRR
AT4G31250            TVFLLALTILAVVVLITVFLSVCILSRRQGK
AT4G32000            LLIALIITSSSLGLILVSCLCFWVYWSKKSP
AT4G34440_PERK5      GLIIGVLVGAGLLLLLAVCICICCNRKKKKK
AT4G34500            YLVIAICSVFILLISLLIFLFVCLNRVSRAR
AT4G37250            VIIGIVVGDIAGIGILAVIFLYIYRCKKNKI
AT5G05160            IIAIVVGCSVAVLFLGIVFLVCLVKKTKKEE
AT5G10290[P]         GIIAGVVAGVTVVLFGILLFLFCKDRHKGYR
AT5G11020            LVISLAATFSLVGIILLCSLLYWFCHRRRNL
AT5G14210_LRR-RLK    LIAVIGGAVLVLVFFVLVILLLLCTNRCSSC
AT5G16590_LRR1[P]    AIVGIVIGCFVLLLVLFLIVFCLCRKKKKEQ
AT5G20480_EFR        VSGICIGIASLLLIIIVASLCWFMKRKKKNN
AT5G24010            VWIVVGSVLGGFVFLSLFFLSVLCLCRRKNN
AT5G24100            ILGIAISVCFVIFFVIAVVIIVCYVKRQRKS
AT5G28680_ANX2       ITAFVIGSAGGVAAVLFCALCFTMYQRKRKF
AT5G35370            IALVLLPCSGFFLLIALGLLWWRRCAVMRYS
AT5G42120_LECRKS6    WSLLPGLAAIVILVAFIVFSLICGKKRISEE
AT5G46330_FLS2[*]    ILIIILGSAAALLLVLLLVLILTCCKKKEKKI
AT5G54380_THE1[P]    IIGSLVGAVTLILLIAVCCYCCLVASRKQRS
AT5G58300[P]         IIPIAAGGAALLLLITVIILCCCIKKKDKRE
AT5G61480_TDR        GAIVWILAAAIGVGFFVLVAATRCFQKSYGN
AT5G65240            IIAGVVSGIAVILLGFFFFFFCKDKHKGYKR
```

**Figure S2:** Alignment of transmembrane domains (TM) for all Arabidopsis RLK proteins that have cysteines within -1,+5 of the end of the TM domain. The alignment is based on the TM domain only (green) and shows the TM domain plus 10 residues after end of the TM domain. The residues within -1+5 of the TM domain end are enclosed by a red box and the cysteine residues that are found within this region are highlighted in orange.
