## Supplementary figures and images for "Juxta-membrane S-acylation of plant receptor-like kinases – fortuitous or functional?"

### Supplemental figure 3

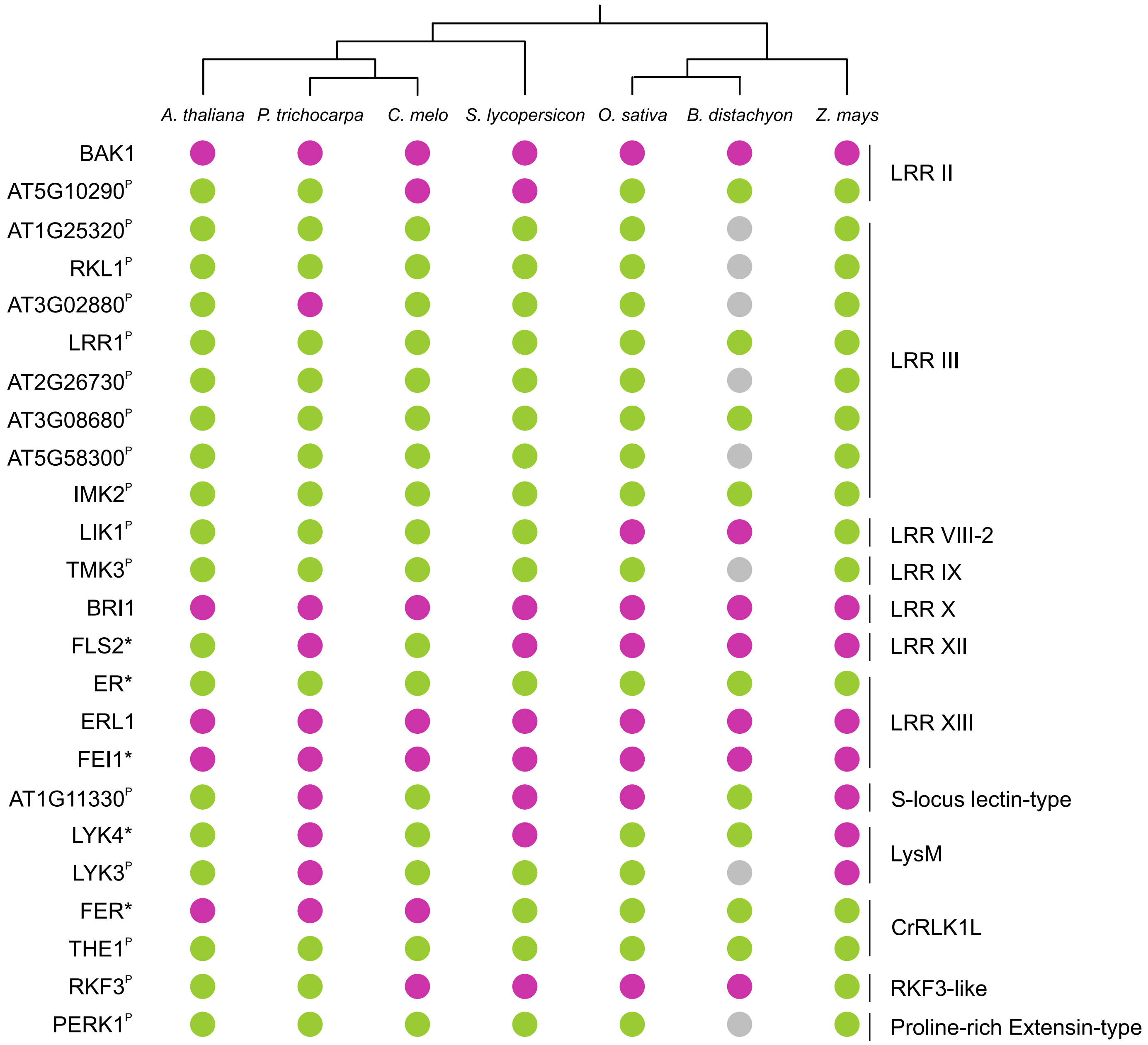
